# A role of DAO1 in oxidation of IAA amino acid conjugates revealed through metabolite, high throughput transcript and protein level analysis

**DOI:** 10.1101/2020.10.24.353276

**Authors:** Müller Karel, Dobrev I. Petre, Pěnčík Aleš, Hošek Petr, Vondráková Zuzana, Filepová Roberta, Malínská Kateřina, Helusová Lenka, Moravec Tomáš, Katarzyna Retzer, Harant Karel, Novák Ondřej, Hoyerová Klára, Petrášek Jan

## Abstract

Auxin metabolism is, together with auxin transport, a key determinant of auxin signalling output in plant cells, yet details on the underlying mechanisms and factors involved are still largely unknown. Processes involved in the auxin metabolism are subject to regulation based on numerous signals, including auxin concentration itself. Altered auxin availability and the subsequent changes of auxin metabolite profiles can therefore elucidate the function and regulatory role of individual elements in the auxin metabolic machinery.

After analysing auxin metabolism in auxin dependent tobacco BY-2 cell line grown in presence or absence of synthetic auxin 2,4-D we found that both conditions were similarly characterized by very low levels of endogenous indole-3-acetic acid (IAA) and its metabolites. However, metabolic profiling after exogenous application of IAA uncovered that the concentration of N-(2-oxindole-3-acetyl)-L-aspartic acid (oxIAA-Asp), the most abundantly formed auxin metabolite in the control culture, dramatically decreased in auxin-starved conditions. To describe the molecular mechanism behind this regulation, we analysed transcriptome and proteome changes caused by auxin starvation. While no changes in the expression of auxin biosynthetic machinery were observed, many genes related to auxin conjugation and degradation showed differential expression. Selected putative auxin glycosylating enzymes as well as members of the Gretchen Hagen 3 gene family involved in auxin amino acid conjugation showed both up- and down-regulation. Contrarily to that, all tobacco homologs of *Arabidopsis thaliana* DIOXYGENASE FOR AUXIN OXIDATION 1 (DAO1), known to be responsible for the formation of oxIAA from IAA, showed significant downregulation at both transcript and protein levels. To validate the role of DAO1 in auxin metabolism, we performed auxin metabolite profiling in BY-2 mutants carrying either siRNA-silenced or CRISPR-Cas9-mutated *Nt*DAO1, as well as in *dao1-1 Arabidopsis thaliana* plants. Both mutants showed not only expectedly lower levels of oxIAA, but also significantly lower abundance of oxidated amino acid conjugates of IAA (oxIAA-Asp). Our results thus represent the first direct evidence on DAO1 activity on IAA amino acid conjugates.

**Statement of significance:** Here we present an analysis of auxin metabolism on metabolite, transcript and protein levels in tobacco BY-2 cell line, collectively identifying oxidation of IAA amino acid conjugates as a new role of DIOXYGENASE FOR AUXIN OXIDATION 1 within an auxin-level-responsive metabolic system.

## Introduction

Auxin is one of the most important plant morphogenic compounds regulating many processes of plant development at several levels (Paque and Weijers, 2016). Concentration of auxin in plants, mainly represented by indole-3-acetic acid (IAA), is tightly regulated through biosynthesis, transport, degradation and conjugation; the last involving both reversible and irreversible pathways. Together, these processes maintain auxin homeostasis which, when disrupted, leads to physiological or morphological changes (Ludwig-mu, 2014; Casanova-Sáez and Voß, 2019). Transport across the plasma membrane, facilitated mostly through activities of PIN and AUX/LAX proteins, is probably the most studied auxin regulatory mechanism. Many detailed studies focused on localization, regulation of activity as well as kinetic parameters have been published in recent years (Reemmer and Murphy, 2014; Adamowski and Friml, 2015; Sauer and Kleine-Vehn, 2019). Contrary to that, a plethora of blank spots remain in auxin metabolism. While several mechanisms of auxin biosynthesis were predicted, only one pathway - that with intermediate indole-3-pyruvate catabolized by members of tryptophan aminotransferase and flavin monooxygenase families - was identified in plants (Mashiguchi et al., 2011). Auxin oxidation is a major way of its degradation but the enzyme involved in a formation of 2-oxindole-3-acetic acid (oxIAA), DIOXYGENASE FOR AUXIN OXIDATION 1 (DAO1), has been identified just recently (Porco *et al.*, 2016; Zhang *et al.*, 2016). Most of the IAA in plant cells is present in conjugated forms, usually in bond with amino acids or sugars even though other forms exist as well. Levels of these metabolites depend on cell type as well as on plant species (Ludwig-Müller, 2011). Ester-linked sugar conjugation in plants is catalyzed by specific uridine-diphosphate glycosyltransferases (UGTs). IAA-specific UGTs have been described for example in *Arabidopsis* (Jackson *et al.*, 2001) and maize (Szerszen *et al.*, 1994), but their wider identification and characterization in other plants is difficult due to the large gene family of UGTs. Amide-bond linked amino acid conjugates (IAA-aa) are products of the activity of GRETCHEN-HAGEN 3 (GH3) gene family of acyl acid-amido synthetases. Some members of this group are known for their promiscuous character due to the range of amino acid donors as well as plant hormone acceptors (Staswick, 2005; Westfall *et al.*, 2016; Aoi *et al.*, 2020). Products of both oxidation and conjugation, for example 2-oxindole-3-acetic acid-glucose (oxIAA-GE) or 2-oxindole-3-acetic acid-amino acid conjugates (oxIAA-AA), have been identified in plants, suggesting close coordination of oxidative and conjugating reactions (Zhang and Peer, 2017). In addition, compartmentalisation plays an important role in auxin metabolism. Several members of PIN proteins and members of PIN-likes (PILS) were localized to the membrane of endoplasmic reticulum (Barbez *et al.*, 2012; Simon *et al.*, 2016) and IAA-aa amidohydrolases ILR1, IAR1 and ILL2 are localized to the ER lumen (Sanchez Carranza *et al.*, 2016).

While a detailed transport-based mechanism of auxin homeostasis in *Nicotiana tabacum* L., cv. Bright Yellow (BY-2) cells was recently revealed, no information is known about the natural metabolism of auxin in BY-2 cells. In the present, work we performed multi-omical characterization of BY-2 cells cultured in the presence (control) or absence (auxin-starved) of synthetic auxin 2,4-dichlorophenoxyacetic acid (2,4-D). Significant qualitative and quantitative changes in auxin metabolic profiles collectively with transcript and protein levels of auxin metabolic machinery suggested a new role of tobacco homolog of DAO1 in oxidation of IAA amino acid conjugates. This novel function of DAO1 was confirmed in mutants of DAO1 in tobacco BY-2 cells and *Arabidopsis* plants.

## Results

### 2,4-D-supplemented and auxin-free-cultured BY-2 cells show the same grow parameters during the first two days of cultivation

Tobacco BY-2 cell line is standardly cultured in the presence of synthetic auxin 2,4-D in the medium. When 2,4-D is not present, auxin deprivation leads to an inhibition of cell division, and induction of cell elongation accompanied by other morphological and biochemical changes (Winicur *et al.*, 1998). In order to select the optimal conditions for analysis of auxin-responsive auxin metabolism, selected growth parameters (i.e. titer, weight and viability of cells) were followed during the growth cycle of BY-2 cells cultured in presence or absence of 2,4-D (Figure 1). In control cells, cell titre and cell weight showed typical exponential growth character with a maximum in the fifth day. Auxin-deprived cells showed no cell division during the seven days of growth cycle (Figure 1A,C). The weight of auxin-free-cultured cells increased slightly with a maximum in the sixth day (approximately 11-fold increase in fresh weight compared to day zero) (Figure 1C). High amount of living cells (approximately 95%) was observed in the 2,4-D-supplemented culture during the whole growth cycle (Figure 1B). Auxin-deprived cells maintained a stable level of living cells during the first three days of cultivation and then dropped to approximately 80% and 60% ratio of viable cells in 4^th^ and 7^th^ day, respectively.

**Figure 1.**
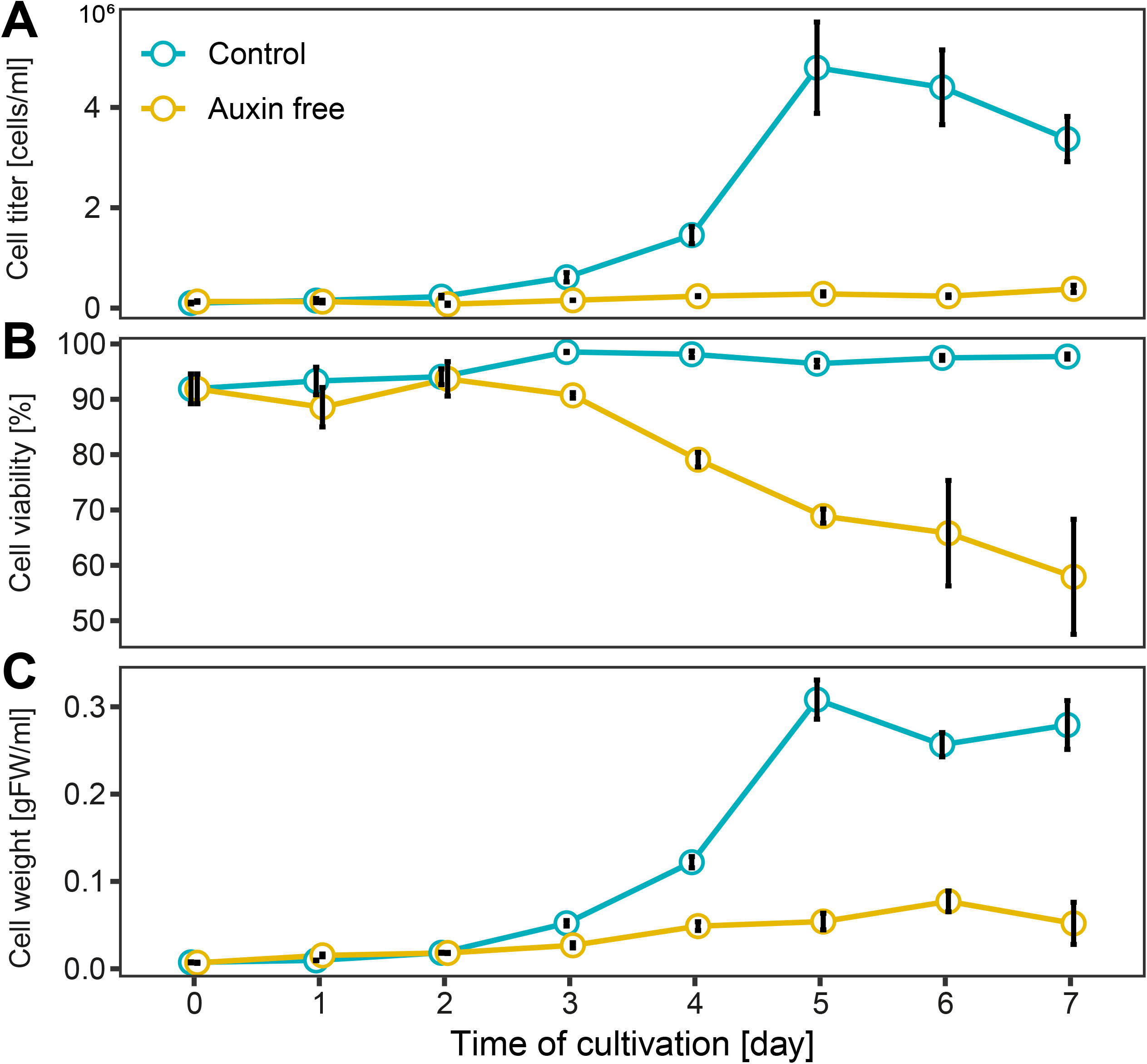
Growth parameters of BY-2 cells in auxin-free medium. Cell titer **(A)**, viability (ratio of living cells) **(B)** and weight **(C)** were monitored during cultivating period of *Nicotiana tabacum* L. cv. Bright Yellow 2 cells cultured in the presence (control) and absence (Auxin free) of synthetic auxin 2, 4-D. Means ± SE are shown (n = 3).

These results show that 2,4-D-supplemented and auxin-deprived BY-2 cells show comparable grow parameters during the first two days of cultivation, therefore we decided to perform analysis of auxin metabolites and auxin metabolism in auxin deprived cells two days after inoculation.

### oxIAA-Asp is the main auxin metabolite in BY-2 cells

Levels of endogenous IAA and its metabolites were determined by mass spectrometry analysis in two days old BY-2 cells cultured in presence (control) or absence (AF) of synthetic auxin 2,4-D (Figure 2A). The major IAA metabolite in both conditions was oxIAA-Asp with levels almost 100-fold higher compared to other metabolites oxIAA, oxIAA-GE and IAA. While the level of IAA was similar in both conditions, the level of oxIAA was 3-fold lower in auxin-starved cells. Contrarily to that, auxin-deprived cells showed slightly higher abundance of oxidized forms of IAA conjugates oxIAA-Asp and oxIAA-GE (1.6- and 1.1-fold, respectively). Levels of IAA-GE, IAA-Asp and IAA-Glu were below the detection limit of the instrument. To exclude the possibility that metabolites are exported out from the cells, we harvested and analyzed the culture media but no measurable levels of either IAA or its metabolites were found.

**Figure 2.**
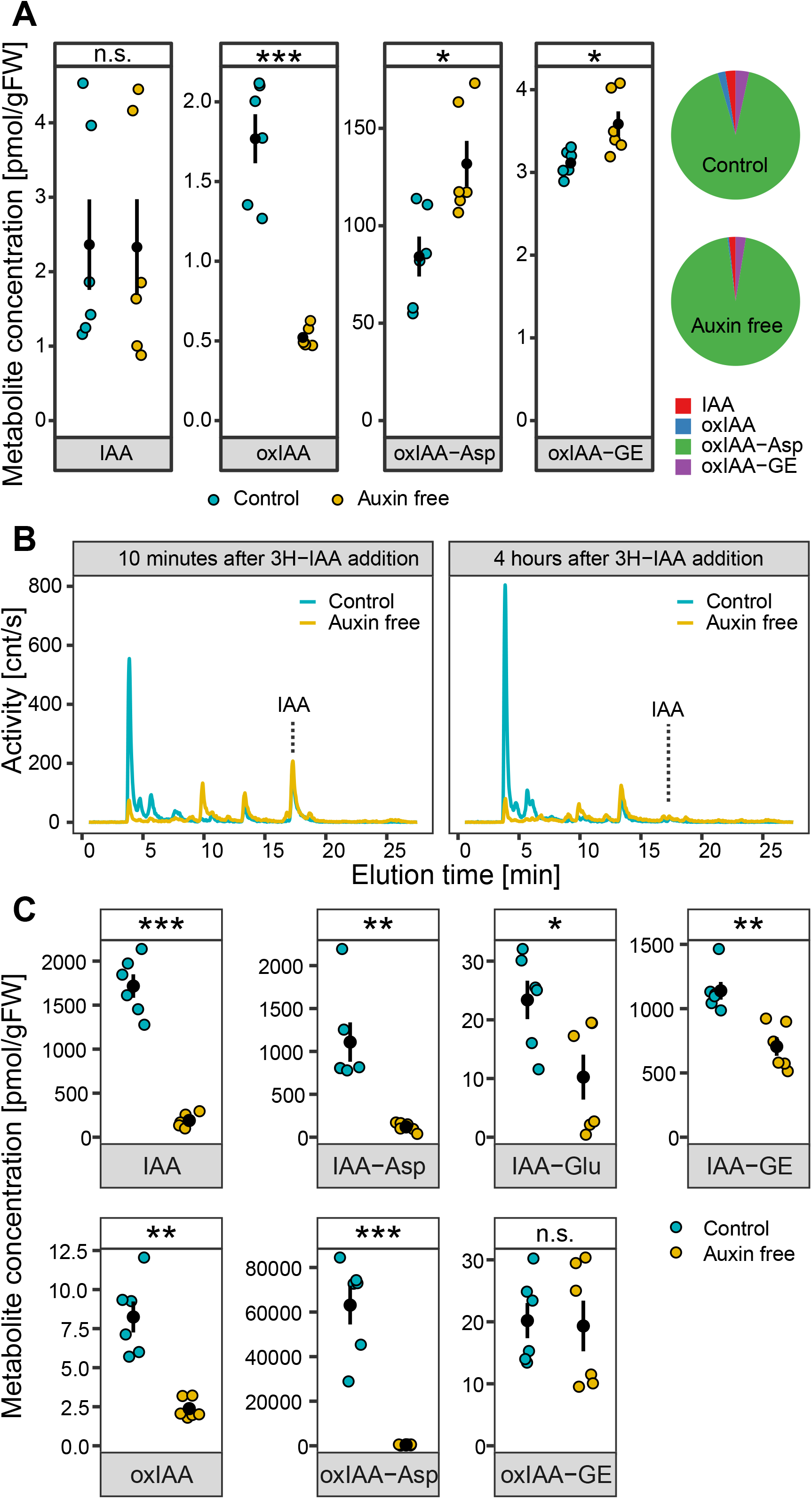
Distinct auxin metabolism in 2, 4-D supplemented and auxin free cultivated BY-2 tobacco cell culture **(A)** The endogenous contents of free IAA and its metabolites were measured in two days old control (2,4-D supplemented) and auxin free cultivated BY-2 cell culture. Data points and error bars indicating mean ± SE are shown. Pie charts represent relative contribution of detected metabolites to the overall content of auxin in cells. **(B)** Auxin metabolite profiling of control and auxin free cultivated cells performed by HPLC chromatography ten minutes and four hours after application of ^3^H-IAA into the media. **(C)** Mass spectrometry based quantification of free IAA and its metabolites in control and auxin free cultivated cells two hours after application of 1μM IAA into the media. Data points and error bars indicating mean ± SE are shown. Student t-test P-values: *P<0.05, **P<0.01, ***P<0.001

### 2,4-D controls the capacity of BY-2 cells to metabolize exogenously applied IAA

Two days old control and auxin-free BY-2 cells were compared in their ability to metabolize exogenously applied IAA. Quantitative and qualitative analyses of IAA and its metabolites were performed by high-performance liquid chromatography (HPLC) and mass spectrometry (MS), respectively. Radioactively labelled IAA and its metabolites were analyzed in cellular extracts 10 minutes and 4 hours after exogenous application of ^3^H-IAA. Figure 2B shows significantly different metabolic profiles between control and auxin-deprived BY-2 cells. The dominant metabolite in control cells, represented by a peak eluted in the 4^th^ minute, was present with a very low abundance in AF sample. ^3^H-IAA, represented by a peak eluted in the 17^th^ minute, was completely metabolized in both conditions 4 hours after its application. However, all IAA metabolites were generally present at much lower levels in auxin-deprived cells.

Mass spectrometry was used to validate the qualitative results of HPLC analysis (Figure 2C). Levels of IAA and its metabolites were measured in BY-2 cells 2 hours after application of IAA into the media. Higher abundance of all metabolites except for oxIAA-GE was detected in control cells compared to auxin-deprived cells. The most striking difference was observed in case of oxIAA-Asp, which was the dominant metabolite in control (150-fold higher abundance compared to AF). In conclusion, we found significant quantitative and qualitative auxin-dependent changes in the capacity of BY-2 cells to metabolize exogenously applied IAA.

### High-throughput analysis of RNA and protein levels in control and auxin-free-cultured BY-2 cells

To understand the molecular background responsible for auxin-dependent differences in IAA metabolic capacity of BY-2 cells, transcriptomic and proteomic analyses were carried out. RNA-seq was performed for 7 control and 3 auxin-free-cultured BY-2 cell culture samples harvested two days after inoculation. Quality of RNA and total number of reads per biological replicate are summarized in Table S1.

Reads were quality-filtered and mapped against *Nicotiana tabacum* v1.0 cDNA reference dataset (Edwards *et al.*, 2017) using the software salmon (Patro *et al.*, 2017). Quality of the analysis was assessed by principal component analysis and hierarchical clustering (Figures 3A, S1). Transcripts with TPM value higher than 0.5 in either control or AF were considered as detected (totally 40285 transcripts out of 69500 in the reference dataset). DESeq2 (version 1.18.0, (Kumagai-Sano *et al.*, 2007)(Kumagai-Sano *et al.*, 2007)(Kumagai-Sano *et al.*, 2007)(Kumagai-Sano *et al.*, 2007)Love *et al.*, 2014) and Sleuth (version 0.29.0, Pimentel *et al.*, 2017) R packages were used for identification of differentially expressed genes (DEGs). 23348 transcripts were agreed upon by both algorithms as being significantly differently expressed (q-value <= 0.05). Of these transcripts, 8770 and 7598 were considered as up- and down-regulated, respectively (fold-change >= 2, summarized in Table S2). In brief, members of the extensin gene family and stress-related haem peroxidases were among the most up-regulated genes, while members of AUX/IAA and Gretchen Hagen 3 families belonged to the most down-regulated genes in auxin-free cultured BY-2 cells.

**Figure 3.**
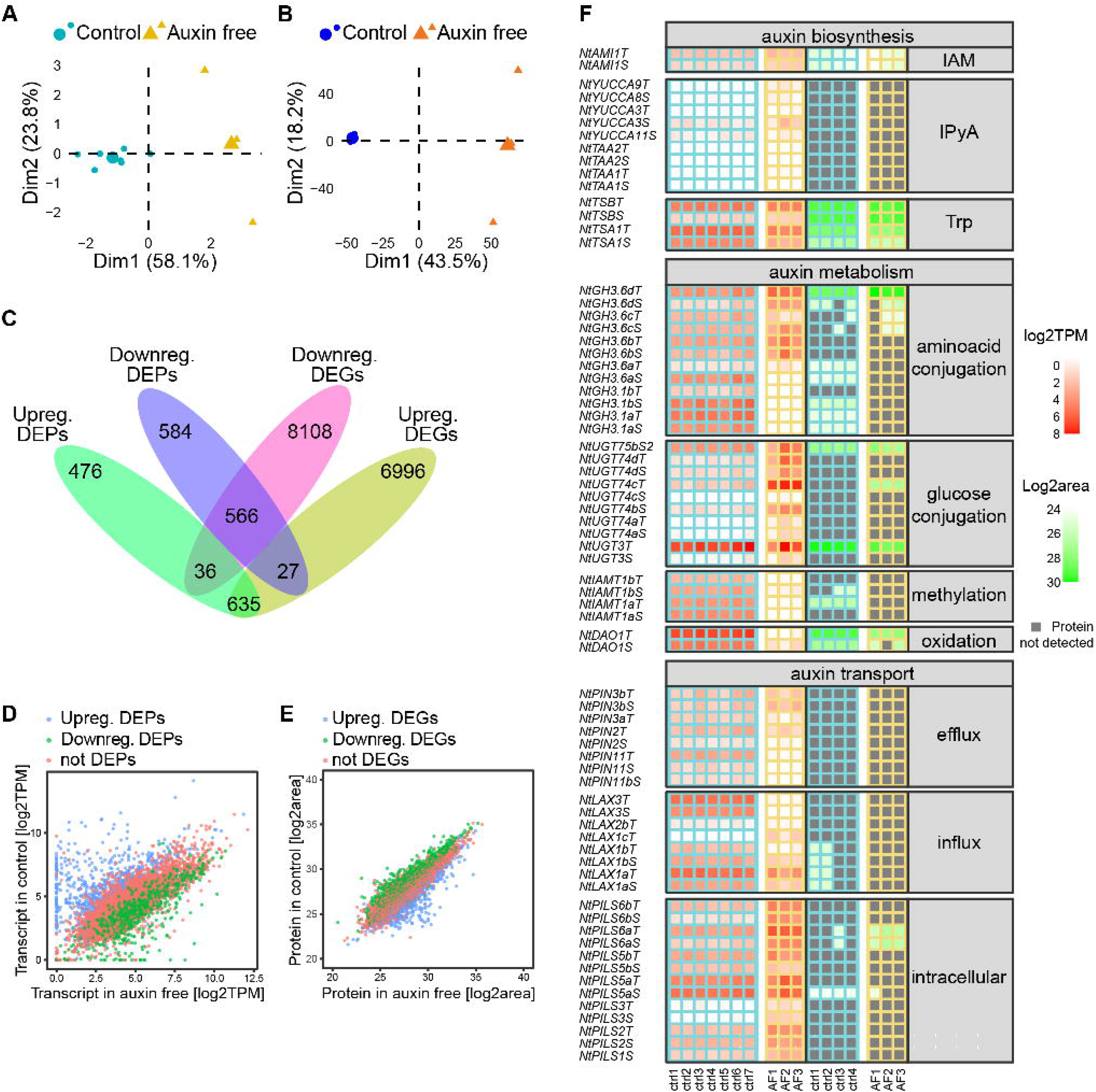
Transcriptomic and proteomic characterization of 2,4-D supplemented and auxin starving tobacco BY-2 cell culture. PCA of RNAseq **(A)** and mass spectrometry defined proteome **(B)** of two days old BY-2 cell culture. RNAseq was performed for 7 control (2,4-D supplemented) and 3 auxin free samples. Proteomic analysis was done using 4 control and 3 auxin free samples. Venn diagram **(C)** and scatter plots **(D, E)** summarize differentially expressed genes and proteins (DEGs and DEPs) and their overlap in both methods. Plot **(D)** relates transcription abundance (TPM) in control and auxin free samples, colour of dots represents results of statistics for particular protein in proteome analysis. Plot **(E)** relates protein abundance (log2 transformed area of peaks), colour of dots represents result of statistics in RNAseq experiment. In **(F)**, heat map representing transcription and protein levels of selected genes involved in the regulation of auxin activity (biosynthesis, metabolism and transport) is shown.

To address the biological processes encoded within the differentially expressed genes, we used the GO statistical overrepresentation test of PantherDB (Mi *et al.*, 2019). In summary, processes associated with succinate transmembrane transport, calcium-dependent signalling, beta-glucan biosynthesis, plant-type cell wall biogenesis and cytoskeleton-dependent cytokinesis were enriched in genes up-regulated after auxin removal. Processes associated with mitochondrial translation and membrane organization, protein synthesis (ribosome biogenesis, translation, protein folding and methylation) and cell division (DNA replication, chromosome segregation and organization, DNA repair and recombination) were enriched in down-regulated set of transcripts. Complete results of PANTHER Overrepresentation tests are shown in Tables S3 and S4)

Proteomic characterization of BY-2 cells in auxin-supplemented and auxin-free conditions was done by label-free quantitative LC-MS/MS analysis. Quality control of the analysis was done by principal component analysis and hierarchical clustering (Figures 3B, S2). In total, 6547 proteins were identified in 2-day-old BY-2 cells (q-value <= 0.05). 2324 proteins were determined as differentially expressed (DEPs), q-value <= 0.05, fold-change >= 2) with 1147 and 1177 showing up- and down-regulated pattern, respectively. Peroxidases, extensins, annexin, methyltransferase and proteinase inhibitor represent the most up-regulated proteins, whereas histone H2A, phospholipase D, helicase, several members of ribosomal proteins or glyceraldehyde-3-dehydrogenase belong to the most down-regulated ones. Combining the transcriptomic and proteomic data we found 566 downregulated and 635 upregulated genes at both their transcript and protein levels. 37 genes were upregulated at the protein level and downregulated at transcript levels and 27 genes showed the opposite pattern (summarized in Venn diagram in Figure 3C). Visualization of correlation of DEGs and DEPs is shown as scatter plots in Figures 3D, E. Altogether, our data provide the first high-throughput analysis of the BY-2 cell line with very robust characterization of transcript and protein levels.

### Distinct regulation of genes involved in auxin metabolism

The RNA-seq and proteomic data were filtered to select genes related to auxin metabolism (biosynthesis, conjugation and degradation) and transport to better understand the molecular background for different auxin metabolic capacity of auxin-supplemented and auxin-deprived BY-2 cells. Transcription of selected genes was validated by RT-qPCR (Figure S3).

Auxin biosynthesis was represented by homologs of tryptophan aminotransferases (TAAs), flavin-binding monooxygenases (YUCCAs), amidase1 (AMI1), aldehyde oxidase 1 (AAO1) and tryptophane synthases (TSA and TSB). Genes coding for enzymes involved in indole-3-pyruvic acid pathway (TAAs and YUCCAs) showed very little transcription in both, control and auxin-free conditions. The proteins coded by these genes were not detected at all. Putative indole-3-acetamide pathway was represented by AMI1, which was expressed at similar levels in both conditions. Genes involved in tryptophane synthesis show higher expression in control cells (Figure 3F).

Next, we focused on genes related to auxin conjugation and degradation, in particular members of *GRETCHEN HAGEN 3* gene family involved in IAA amino acid conjugation, *DIOXYGENASE FOR AUXIN OXIDATION 1 (DAO1)* involved in oxidation of IAA, IAA carboxymethyltransferase (*IAMT*) and selected members of UDP-glucosyl transferases (*UGTs*). Since plant genomes harbor many genes coding for UGTs, we selected homologs of *AtUGT84B1* (AT2G23260), which is responsible for IAA glycosylation in *Arabidopsis thaliana*. *NtUGT3* was added to the UGT selection for its high expression in BY-2 cell culture. Expressions of *NtUGT3*, several *GH3* members, as well as tobacco homologs of *IAMT* and *DAO1* were significantly lower in auxin-free cultured BY-2 cells compared to control. Surprisingly, *GH3.6b, c* and *d* isoforms of the *GRETCHEN HAGEN 3* family show no change upon auxin removal or even increase in the expression level. Furthermore, *NtGH3.6dT* showed the highest expression among GH3 members at the protein level in both control and auxin-free culture BY-2 cells (Figure 3F).

Genes coding for auxin transporters were represented by genes from the auxin influx carrier family (*AUX/LAXes*), auxin efflux carriers (*PINs*) and putative endomembrane auxin transporters (*PILSes*). The highest expression at both transcript and protein levels was found in case of PILS5 and PILS6 homologs and their abundance was significantly higher in auxin-free cultured cells. Contrarily to that, genes of the PIN family showed generally lower transcript levels and all expressed members except for PIN3b were downregulated in auxin-free conditions. The *LAX* gene family was represented by the dominant transcripts of *LAX1a* and *LAX3*. Auxin deprivation caused downregulation of most tobacco *AUX/LAX* genes and induction of expression of *LAX1c*, confirming the results published in Müller *et al.*, 2019.

To conclude, here we show a multi-omical comparison of tobacco BY-2 cells in auxin-supplemented and auxin-free conditions. No significant changes in auxin biosynthesis were detected. However, multiple genes coding for auxin-metabolizing enzymes showed changes at transcript and/or protein levels. Amino acid conjugating pathway, represented by the Gretchen Hagen 3 gene family, showed both up and down regulation. Genes from the family of UDP-glucosyltransferases, coding for putative IAA glycosylating enzymes, were generally more expressed in auxin-deprived cells. However, NtUGT3, the most abundant UGT member at the transcript and protein level, was less expressed in auxin-free-cultured cells, suggesting its role in auxin metabolism. The homologs of DIOXYGENASE FOR AUXIN OXIDATION (DAO1) showed significant downregulation upon auxin removal at both transcript and protein levels. We propose that the different expression of DAO1 is a crucial factor for significantly different metabolic profiles, in particular for the presence and absence of oxIAA-Asp between the control and auxin-free-cultured BY-2 cells. To prove our hypothesis, we prepared BY-2 mutant lines with altered activity of DAO1 protein.

### IAA-Asp is the main IAA metabolite in DAO1 knockout mutant of BY-2 cell culture

BY-2 line with knocked-down DAO1 activity (siDAO1) was prepared by β-estradiol-inducible expression of DAO1 specific siRNA. Reduced expression of DAO1 in β-estradiol-treated cells (approximately 2- and 3-fold lower compared to WT and DMSO-treated control, respectively) was verified by qPCR (Figure S4). BY-2 line with knocked-out DAO1 activity (crisprDAO1) was prepared using CRISPR-Cas9-based genome editing, targeting two sites in the genomic sequences of NtDAO1S and NtDAO1T. Verification of the crisprDAO1 line was done by PCR amplification of the CRISPR-targeted DAO1 genomic sequence followed by cloning of the product into pJET1.2 vector and Sanger sequencing (Table S5).

Transgenic lines siDAO1 and crisprDAO1 were tested for their capacity to metabolize exogenous IAA. Concentrations of intracellular IAA and its metabolites were measured by mass spectrometry two hours after application of 1µM IAA. Figure 4A shows that crisprDAO1 line metabolized IAA primarily into amino acid conjugates IAA-Asp and IAA-Glu (76- and 22-fold increase compared to WT control), whereas the abundance of oxidized forms represented by oxIAA, oxIAA-Asp and oxIAA-GE was in comparison to WT BY-2 cells 2.5-, 120- and 3-fold lower, respectively. To support the effect of modulated activity of DAO1, EST-siDAO1 cells in comparison to DMSO-siDAO1 control showed the concentrations of IAA-Asp and IAA-Glu 10- and 6-fold higher, respectively, while all other metabolites showed similar levels. These measurements support the role of DAO1 in oxidation of IAA to form oxIAA, but also present its new role in the oxidation of IAA-Asp to form oxIAA-Asp.

**Figure 4.**
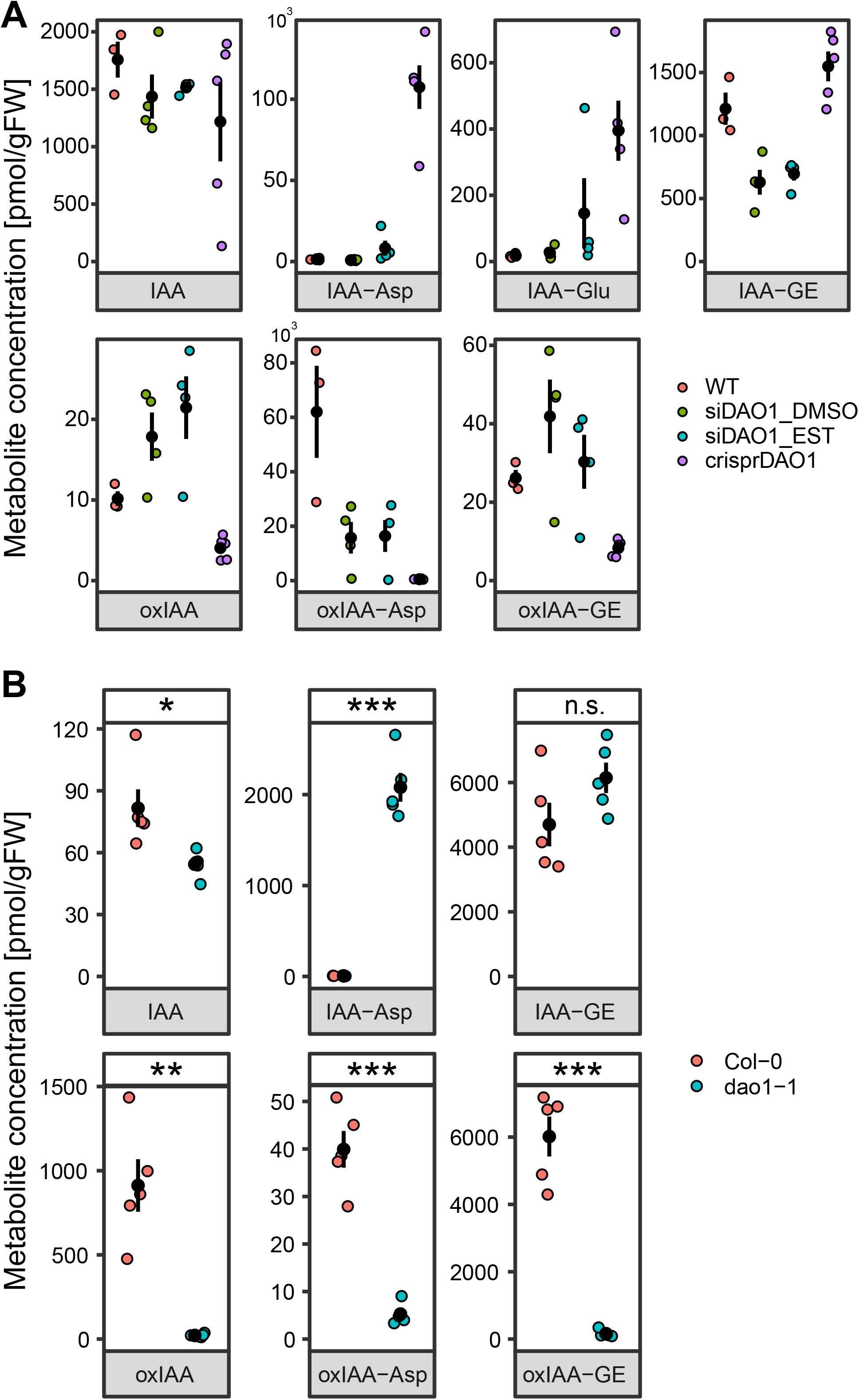
Activity of DAO1 correlates with the levels of oxIAA as well as of oxidized conjugated forms of IAA oxIAA-Asp and oxIAA-GE. Mass spectrometry based quantification of IAA and its metabolites in BY-2 cell lines **(A)** and seedlings of *Arabidopsis thaliana* **(B)**. In **(A)**, metabolites were measured in control BY-2 cell culture (WT), DMSO-treated β-estradiol-inducible DAO1-silencing cell line (siDAO1_DMSO), β-estradiol-induced DAO1-silencing cell line (siDAO1_EST) and CRISPR-Cas9 mutated DAO1 cell line (crisprDAO1) 2 hours after application of 1μM IAA into the media. In **(B)**, IAA and its metabolites were measured in Col-0 and dao1-1 KO mutants. Data points and error bars indicating mean ± SE are shown. Student t-test P-values: *P<0.05, **P<0.01, ***P<0.001

To verify the role of DAO1 in *Arabidopsis thaliana* plants, we performed profiling of endogenous IAA metabolites in *dao1-1* mutant. There, significantly lower levels of oxIAA and oxIAA-GE were accompanied by significantly lower level of oxIAA-Asp metabolite (approximately 10-fold lower in dao1-1 compared to Columbia-0 plants)(Figure 4B). Based on previously reported auxin metabolite analysis in tobacco and *Arabidopsis* mutants, we conclude that the role of DAO1 in auxin metabolism is broader than previously believed because of its activity on amino acid conjugates discovered here.

### DAO1 is localized to the nucleus and cytosol of tobacco BY-2 cells

To find whether DAO1 is under auxin-dependent posttranslational regulation, transgenic BY-2 lines carrying GFP-translational fusion to the coding sequence of NtDAO1T under native (pDAO1T:DAO1-GFP) or under the strong constitutive promoter G10-90 (G10-90:DAO1-GFP) were prepared. Images of two-day-old BY-2 cells cultured in control or auxin-starvation conditions were obtained by confocal microscopy. In auxin-supplemented conditions, the GFP signal was strong in cytosol and nucleus in G10:90:DAO1-GFP as well as in pDAO1T:DAO1-GFP line (Figure 5A, B). In auxin-free conditions, the signal was weak in pDAO1:DAO1-GFP line, whereas G10-90:DAO1-GFP line showed a GFP signal of an intensity similar to auxin-supplemented conditions (Figure 5A, B). We conclude that the auxin-dependent pattern is largely controlled at the transcription level. No signal was observed in control WT BY-2 line (Figure 5C). Cytosolic and nuclear localization of NtDAO1T was confirmed in leaves of *Nicotiana benthamiana* infiltrated by *Agrobacterium thumefaciens* carrying 35S:DAO1-GFP gene construct (Figure 5D)

**Figure 5.**
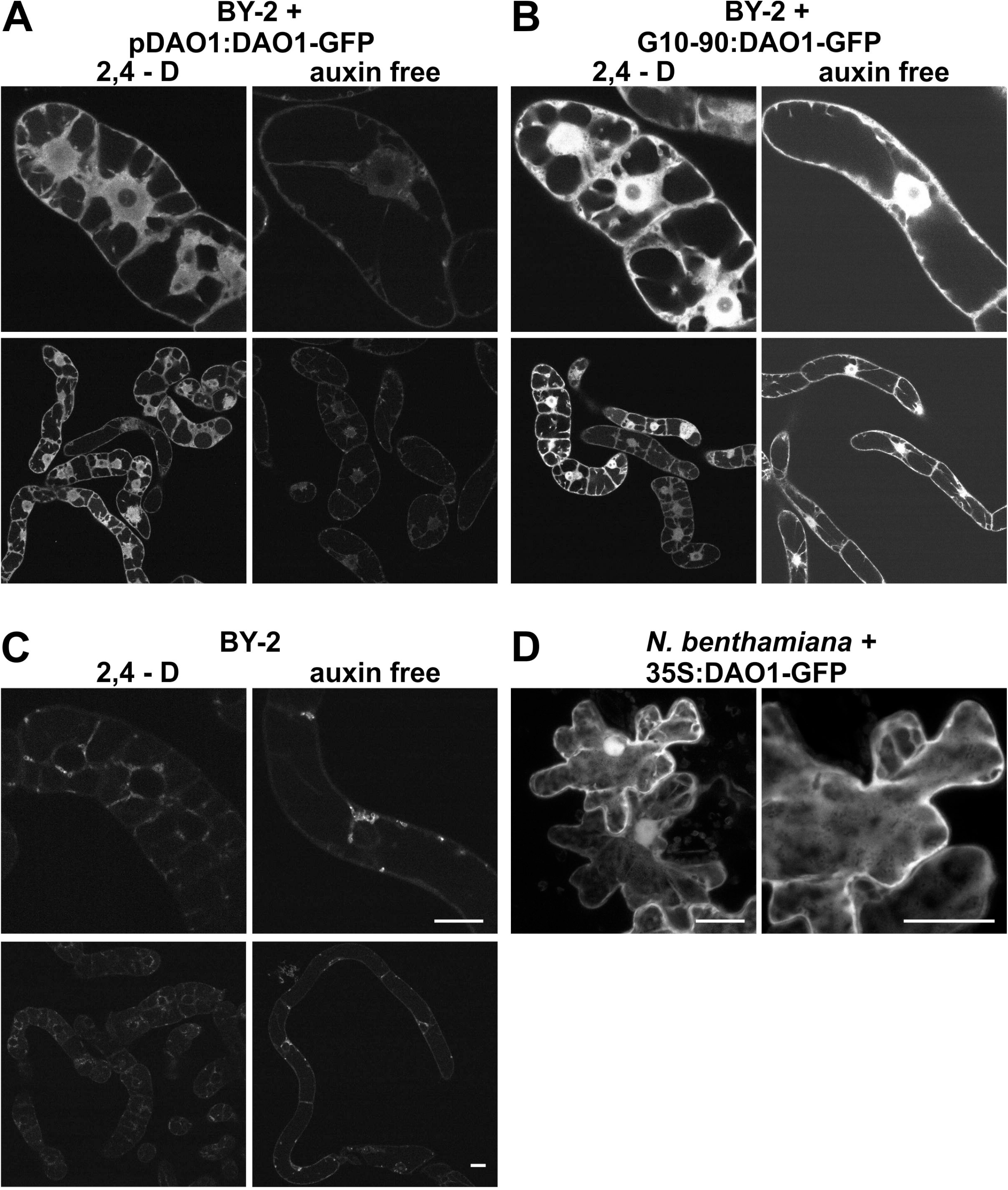
Subcellular localization of DAO1-GFP in BY-2 cells and in tobacco leaves. **(A-C)** Confocal images of 2 days old *Nicotiana tabacum* L. cv. Bright Yellow 2 transgenic pDAO1T:DAO1-GFP **(A)**, G10-90:DAO1-GFP **(B)** lines and wild-type BY-2 line **(C)** in 2,4-D-supplemented or auxin free conditions are shown. DAO1-GFP regardless its promoter is localized in cytosol and nucleus without nucleolus in auxin supplemented conditions (**A, B** – 2,4-D). In auxin free conditions, DAO1 promoter driven DAO1-GFP signal is very week (**A** - auxin free). Individual optical sections are presented. Identical scanning parameters were used during acquisition so that fluorescence intensities should reflect expression levels in cell lines presented in A-C. **(D)** Agrobacterium infiltrated *Nicotiana benthamiana* leaves expressing 35S:DAO1-GFP. Maximum intensity projection of confocal z-stacks. Bars 20 μm.

## Methods

### Plant material and chemicals

Tobacco cell line BY-2 (*Nicotiana tabacum* L. cv. Bright Yellow 2) was maintained in liquid Murashige and Skoog (MS) medium (3% sucrose (w/v), 4.3 g/l MS salts, 100 mg/l myo-inositol, 1mg/l thiamin, 0.2 mg/l 2,4-dichlorophenoxyacetic acid (2,4-D), and 200 mg/l KH_2_PO_4_; pH 5.8), in the dark, at 27°C, under continuous shaking (150 rpm; orbital diameter 30 mm) and sub-cultured every 7 days. Auxin-free-cultured cells were inoculated to medium lacking 2,4-D. Transgenic cells and calli were maintained on the same medium supplemented with 100 μg/ml cefotaxime and 20 μg/ml hygromycin or 50 μg/ml kanamycin. The Arabidopsis ecotype Columbia and *dao1* T-DNA insertion mutants (obtained from Nottingham Arabidopsis Stock Centre) were grown on Murashige and Skoog square agar plates (4.4 g/l Murashige and Skoog, 10 g/l sucrose and 10 g/l plant agar, pH 5.7). Plates were stratified for 4 d and placed vertically in long-day conditions (16 h light/8 h dark) at 22°C. Whole seedlings were collected in five replicates and weighed (approx. 5 mg per replicate), immediately frozen in liquid nitrogen and stored at −80 °C until extraction.

Unless otherwise stated, all chemicals and kits were obtained from Sigma-Aldrich Inc.

### RNA isolation and sequencing analysis

Total RNA was isolated using RNeasy Plant Mini kit (Qiagen) from 50-100 mg of BY-2 cells and treated with DNA-Free kit (Thermo Fischer Scientific). RNA purity, concentration and integrity was evaluated on 0.8% agarose gels and by the RNA Nano 6000 Assay Kit using Bioanalyzer instrument (Agilent Technologies). For RNA-seq analysis, approximately 5 μg of RNA isolated from seven control and three auxin-free replicates were submitted for the service procedure provided by GATC-Biotech company. The analysis resulted in at least 20 million 50 bps long reads. Rough reads were quality-filtered using FastQC and Trimmomatic (parameters: leading:3, trailing:3, slidingwindow:4:20, minlen:40). Transcript abundances (transcripts per million – TPM) were determined using Salmon (Patro *et al.*, 2017) with parameters –seqBias, --gcBias, --numBootstraps 30. Index was built from Nicotiana tabacum v1.0 cDNA dataset (Edwards *et al.*, 2017). Differentially expressed genes were selected using sleuth and DeSeq2 packages in R (Pimentel *et al.*, 2017; Love *et al.*, 2014). Transcripts with q-value <= 0.05 determined by both software and log2 fold change >=1 (upregulated) or <= −1 (downregulated) were considered to be significantly differentially expressed. Gene ontology analysis (statistical overrepresentation test) was done using Panther Classification System (Mi *et al.*, 2019).

### Proteomic analysis

Plant cell callus was lysed with combination of sonication and high temperature detergent-assisted lysis. Proteins were trypsin-digested and nanoflow liquid chromatography was used for separation of the resulting peptides. Next, analysis of the samples by tandem mass spectrometry (nLC-MS2) on a Thermo Orbitrap Fusion (q-OT-qIT, Thermo Scientific) instrument was performed as described elsewhere (Erban *et al.*, 2019). The mass spectrometry data were analysed and quantified using MaxQuant software (version 1.5.3.8) (Cox *et al.*, 2014) with false discovery rate set to 1% for both proteins and peptides and we specified a minimum length of seven amino acids. The Andromeda search engine was used for the MS/MS spectra search against the tobacco protein dataset (Edwards *et al.*, 2017), then normalized intensity values were further processed using the Perseus software (Tyanova *et al.*, 2016).

### Quantitative real time PCR

Approximately 1 μg of DNAse-treated RNA was reverse-transcribed using M-MLV reverse transcriptase, RNase H-, point mutant (Promega). Quantitative real-time PCR was performed using GoTaq qPCR Master Mix (Promega) at 58oC annealing temperature on LightCycler480 instrument (Roche). PCR efficiency was estimated using serial dilution of template cDNA. Tobacco aminopeptidase-like, fructokinase2 and ribokinase-like genes (GenBank accession numbers XM_016640300, KJ683760, XR_001650512, respectively) were used as references in validation of control vs auxin-free RNA-seq data for their stable transcriptions in BY-2 line in our experimental setup. Relative transcript abundancies were then calculated using equation:

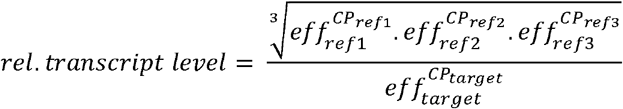

Where eff_ref_ and eff_target_ stand for qPCR efficiency of reference and target genes, respectively, and CP_ref_ and CP_target_ stand for crossing point of reference and targets genes. Positive transcript levels and the quality of PCR products were verified by melting curve analysis. Verification of transgenic lines by RT-qPCR was performed after induction of gene expression with 1 μM β-estradiol. DMSO-treated cells were used as a non-induced control. Sequence of specific primers is shown in Table S5.

### IAA metabolic profiling

Two-day-old BY-2 cells were used for IAA metabolite profiling in experiments aimed at endogenous hormone determination or feeding experiments with free IAA or radioactively labelled IAA (^3^H-IAA). In auxin feeding experiments, control and auxin-free-cultured BY-2 cells were incubated with 20 nM ^3^H-IAA for 4 hours or 1 μM IAA for 2 hours. Auxin metabolites were extracted and purified according to Dobrev and Kamínek (2002) and Dobrev *et al.* (2017)(Ivanov Dobrev and Kamínek, 2002; Dobrev *et al.*, 2017), followed by quantitative analysis on LCMS (endogenous hormone analysis, IAA-feeding experiments) or by the separation on HPLC and detection on Ramona 2000 flow-through radioactivity detector (^3^H-IAA feeding experiments). The metabolites were separated on Kinetex EVO C18 column (2.6 μm, 150 × 2.1 mm, Phenomenex). Mobile phases consisted of A) 5 mM ammonium acetate in water, and B) 95/5 acetonitrile/water (v/v). The following gradient programme was applied: 5% B in 0 min, 7% B in 0.1 to 5 min, 10 to 35 % in 5.1 to 12 min, 100 % B at 13 to 14 min, and 5% B at 14.1 min. LCMS system consisted of UHPLC 1290 Infinity II (Agilent Technologies) coupled to 6495 Triple Quadrupole mass spectrometer (Agilent Technologies). MS analysis was done in MRM mode, using isotope dilution method. Data acquisition and processing was done with Mass Hunter software version B.08 (Agilent Technologies).

Measurement of auxin metabolites in Arabidopsis plants was done following the methods described by Pěnčík *et al.* (2018) (Pěnčík *et al.*, 2018). Samples were extracted with 1 ml of 50 mM phosphate buffer (pH 7.0) containing 0.1% sodium diethyldithiocarbamate and a mixture of stable-isotope-labelled internal standards. 200 μl portion of extract was acidified with 1M HCl to pH 2.7 and purified by in-tip micro solid phase extraction (in-tip μSPE). After evaporation under reduced pressure, samples were analysed using the HPLC system 1260 Infinity II (Agilent Technologies) equipped with Kinetex C18 column (50 mmx2.1 mm, 1.7 μm; Phenomenex) and linked to 6495 Triple Quad detector (Agilent Technologies). Auxin levels were quantified using stable-isotope-labelled internal standards as a reference.

### Preparation of constructs and transformation of BY-2 cells

All vectors for transformation were prepared using GoldenBraid 2.0 cloning system (Sarrion-Perdigones *et al.*, 2013). Basic DNA parts were selected from GoldenBraid 2.0 kit from Diego Orzaez (Addgene kit # 1000000076) or MoClo Toolkit, from Sylvestre Marillonnet (Addgene kit # 1000000044). Specific parts (target gRNA or siRNA sequence) were prepared in the Laboratory of Hormonal Regulations in Plants IEB or Laboratory of Virology IEB, Prague. The cloning was performed using Type IIS restriction enzymes *BsmBI*, *BsaI* (TermoFischer Scientific) and T4 ligase (Promega). For CRISPS-Cas9 editing of NtDAO1 enzymes, vector containing 35S:hCas9:NosT transcriptional unit (TU) and two guideRNAs (guide1: 5’-AGACCCAAAGCCTCATAAAGTGG-3’, guide2: 5’-CTACAGCTTCTGGAGTGAAGTGG-3’) TUs was used. Transformation of BY-2 cells was done by co-cultivation with *Agrobacterium thumefaciens* strain GV2260 (An *et al.*, 1985). Transgenic lines were harvested after 4 weeks, cultured on solid media with kanamycin and tested for the presence of CRISPR-Cas9 activity. Suspension cultures were established from selected lines validated for NtDAO1 mutation rates.

For inducible silencing of *NtDAO1* transcription, β-estradiol-inducible expression (Zuo *et al.*, 2000) of *NtDAO1*-specific (GenBank ID XM_016591349, region 549-785) hairpin RNA gene construct was prepared. BY-2 cell transformation was performed as described above. 10 independent lines were tested by RT-qPCR for the expression of *NtDAO1*. For the experiment of auxin metabolic profile, line with comparable and stable downregulation of the target was used.

### Light microscopy

For in vivo microscopy, a Zeiss LSM 880 inverted confocal laser scanning microscope with a 40× C-Apochromat objective (NA = 1.2 W) was used (http://www.zeiss.com/). GFP fluorescence (excitation 488 nm, emission 489–540 nm) was detected either in single images or in z-stacks. Maximum intensity projection of z-stacks was created using ZEN black software (Zeiss).

## Discussion

Concentration and activity of auxins are key factors determining many processes in plant growth and development. Several mechanisms of regulation auxin activity were described, including transport (cellular influx, efflux and intracellular translocation across endomembranes), reversible and irreversible conjugation or degradation via oxidative pathways. Tobacco BY-2 cell lines have been frequently used for providing a proof of function and for unraveling the molecular background of various proposed regulatory mechanisms, for example during the discovery of the PIN family of auxin efflux carriers (Petrášek *et al.*, 2006) or the effect of altered expression of putative auxin endomembrane transporters on auxin metabolism (Mravec *et al.*, 2009; Barbez *et al.*, 2012). Despite these studies, information concerning the ability of BY-2 cells to regulate auxin activity was very limited. Recently, we revealed a homeostatic mechanism in BY-2 cells maintaining auxin concentration through transcriptional regulation of auxin efflux and influx carriers (Müller *et al.*, 2019). However, insights into the natural capacity of BY-2 cells to metabolize IAA are still largely limited. Hosek *et al.* reported rapid metabolism of synthetic auxin NAA, while 2,4-D, which is a mandatory component of cultivating media of BY-2 cells, is metabolized only very weakly (Hošek *et al.*, 2012).

Here we show that BY-2 cells maintain very low levels of IAA and its metabolites, and that auxin deprivation does not lead to increased IAA biosynthesis. Levels of endogenous IAA in *Arabidopsis* seedlings varied between 20-40 ng/gFW, while BY-2 cells maintain about 0.5 ng/gFW. A major IAA metabolite in BY-2 cells, oxIAA-Asp was present at similar levels in BY-2 cells and *Arabidopsis* seedlings (approximately 30–60 ng/gFW), while IAA-GE, the dominant metabolite in *Arabidopsis* was not detected in BY-2 cells. This suggest an independence of BY-2 cells of IAA achieved through the fact that the auxin signal is secured through the presence of 2,4-D in the medium. In 2,4-D-free conditions however, BY-2 cells cannot compensate for the absence of auxin by synthesis of IAA, as indicated by the unchanged levels of IAA in control and auxin-deprived cells (Figure 2A), and also by no changes in the transcription of the auxin biosynthetic apparatus (Figure 3F). Our observation of phenotype under auxin-deprived conditions revealed similar changes as described in Winicur *et al.*, 1998.

In contrast to IAA biosynthesis, proteins involved in auxin metabolic machinery are present at very high levels in control BY-2 cells (Figure 3F). Exogenously applied IAA is rapidly metabolized mainly to oxIAA-Asp (Figure 2B,C). In auxin-starved cells, the IAA metabolic profile as well as expression of genes involved in auxin metabolism differ significantly. In general, levels of all metabolites except oxIAA-GE were significantly lower in auxin-deprived cells compared to control two hours after application of 1 μM IAA (Figure 2C). Metabolic profile of radioactively labelled IAA revealed several peaks corresponding to unknown metabolites in auxin-deprived cells. Overall lower levels of IAA metabolites in auxin-starved cells are probably caused by lower influx of auxin from the media. Lower transcript and protein levels of several members of AUX/LAX auxin influx carriers (LAX1a, LAX1b, LAX3) in auxin-deprived cells (Figure 3F) confirm this hypothesis, but involvement of other mechanisms (for example rapid efflux of metabolites into the media or the presence of here undetected metabolites) cannot be excluded.

Next, we aimed to explain the most striking difference in IAA metabolites between control and auxin-deprived BY-2 cells – oxIAA-Asp. Here we present, according to our knowledge, the first determination of oxIAA-Asp level in plants. This metabolite is a product of amino acid conjugation and oxidation. GH3 genes, coding for the enzymes catalyzing amino acid conjugation, were identified throughout the plant kingdom and are involved in regulation of many developmental processes (reviewed for example in Staswick, 2005 and Casanova-Sáez and Voß, 2019). Strong induction of expression of many GH3 genes after auxin treatment classified GH3 among early auxin responsive genes, but opposite regulation of some members was reported for example during fruit development (Matsuo *et al.*, 2018). Our transcriptomic and proteomic analysis identified the most abundant GH3 member (NtGH3.6dT, homolog of Arabidopsis GH3.6) to be upregulated in auxin-free conditions. The same regulation pattern was observed for other homologs of AtGH3.6, while homologs of GH3.1 showed significant downregulation after auxin removal (Figure 3F).

Complex regulation of auxin amino acid conjugation cannot explain differences in auxin-dependent IAA metabolism, therefore we focused on the oxidation pathway represented by tobacco homologs of recently identified DIOXYGENASE FOR AUXIN OXIDATION 1 (DAO1)(Porco *et al.*, 2016; Zhang *et al.*, 2016). Auxin metabolic profiling in DAO1 knock-out and knock-down tobacco and Arabidopsis mutants confirmed increased levels of IAA-Asp (as already reported in Mellor *et al.*, 2016). Moreover, significantly lower abundance of oxIAA-Asp in mutants compared to WT controls provides a direct evidence of DAO1-mediated oxidation of IAA-Asp. Our results thus bring a new fact into the biological aspects of DAO1 in auxin metabolism and the mechanism of coordination of auxin oxidative and conjugating pathways.

The sequence of amino acid conjugation and oxidation has not yet been explored but the data indicates that the oxIAA-Asp-forming pathway starts with amino acid conjugation followed by DAO1-mediated oxidation. A recent study describes allosteric regulation of DAO1, during which the enzyme multimerizes in the presence of IAA (Takehara *et al.*, 2020). Dimeric and tetrameric forms showed higher reaction rate and affinity towards IAA compared to DAO1 monomer, however, the *in-vitro* measured IAA oxidation rate of the ‘active’ form was still about 4-fold lower compared to the reaction rate of GH3-catalyzed IAA amino acid conjugation.

While oxIAA-Asp seems to be the most abundant IAA metabolite in tobacco BY-2 cells, *Arabidopsis* plants mainly catabolize IAA into oxIAA-GE formed by glycosylation of oxIAA (Tanaka *et al.*, 2014; Porco *et al.*, 2016). The large content of oxIAA-Asp in IAA-treated tobacco cells as well as that of oxIAA-GE in *Arabidopsis* plants, both formed during DAO1-dependent reactions, indicate a more important role of DAO1 during auxin homeostasis than previously thought.

Furthermore, it is obvious that the final resolution on the balance of oxidation and conjugation rates and ratios can be declared only when all involved pathways and metabolites are known. It has been, for instance, generally accepted that IAA-Asp and IAA-Glu conjugates cannot be hydrolysed and thus does not present storage forms of auxin (LeClere *et al.*, 2002). Our results raise an interesting hypothesis that the rapid DAO1-mediated oxidation depleting the pool of amino acid conjugates might prevent their possible cleavage by hydrolases. Further research into the specificity of DAO1 and IAA metabolic conversion network in *dao1* mutants is needed to prove our hypothesis and bring new insights in this field of auxin homeostasis.

## Conclusions

Here we show a multiomical characterization of tobacco BY-2 cell culture cultivated in the presence of exogenously supplemented synthetic auxin 2,4-D in comparison with cell culture cultivated for 48 hours in auxin-free conditions. We detected no auxin biosynthesis under both conditions. Contrary to that, we proved significant differences in expression of auxin-metabolizing enzymes at transcript and proteins levels. Knockout and knockdown mutants of tobacco DAO1 verified its importance in IAA metabolism and revealed its role in conversion of IAA-Asp into oxIAA-Asp.

## Notes

### Competing Interest Statement

The authors have declared no competing interest.

